# Modulation of the extracellular matrix by *Streptococcus gallolyticus* subsp. *gallolyticus* and importance in cell proliferation

**DOI:** 10.1101/2022.03.14.484142

**Authors:** Ritesh Kumar, John Taylor, Jain Antrix, Sung Yun Jung, Yi Xu

## Abstract

*Streptococcus gallolyticus* subspecies *gallolyticus* (*Sgg*) has a strong clinical association with colorectal cancer (CRC) and actively promotes the development of colon tumors. Previous work showed that this organism stimulates CRC cells proliferation and tumor growth. However, the molecular mechanisms underlying these activities are not well understood. Here, we found that *Sgg* upregulates the expression of several types of collagens in HT29 and HCT116 cells, with type VI collagen (ColVI) being the highest upregulated collagen type. Knockdown of ColVI abolished the ability of *Sgg* to induce cell proliferation and reduced the adherence of *Sgg* to CRC cells. The extracellular matrix (ECM) is an important regulator of cell proliferation. Therefore, we further examined the role of decellularized matrix (dc-matrix), which is free of live bacteria or cells, in *Sgg*-induced cell proliferation. Dc-matrix prepared from *Sgg*-treated cells showed a significantly higher pro-proliferative activity than that from untreated cells or cells treated with the control bacteria. On the other hand, dc-matrix from *Sgg*-treated ColVI knockdown cells showed no difference in the capacity to support cell proliferation compared to that from untreated ColVI knockdown cells, suggesting that the ECM by itself is a mediator of *Sgg*-induced cell proliferation. Furthermore, *Sgg*-treated CRC cells formed significantly larger tumors *in vivo*, whereas *Sgg* treatment had no effect on ColVI knockdown cells, suggesting that ColVI is important for *Sgg* to promote tumor growth *in vivo*. These results highlight a dynamic bidirectional interplay between *Sgg* and the ECM, where *Sgg* upregulates collagen expression. The *Sgg*-modified ECM in turn affects the ability of *Sgg* to adhere to host cells and more importantly, acts as a mediator for *Sgg*-induced CRC cell proliferation. Taken together, our results reveal a novel mechanism in which *Sgg* stimulates CRC proliferation through modulation of the ECM.

**Author Summary:** Colorectal cancer (CRC) is a leading cause of cancer-related death. The development of CRC can be strongly influenced by specific gut microbes. Understanding how gut microbes modulate CRC is critical to developing novel strategies to improve clinical diagnosis and treatment of this disease. *S. gallolyticus* subsp. *gallolyticus* (*Sgg*) has a strong clinical association with CRC and actively promotes the development of colon tumors. However, the mechanisms *Sgg* utilizes to promote tumors are not well understood. Our results showed for the first time a dynamic interplay between *Sgg* and the extracellular matrix. We found that *Sgg* upregulates the expression of collagens which in turn affects the interaction between *Sgg* and CRC cells and mediates CRC cell proliferation. These findings draw attention to a previously unrecognized dynamic bidirectional interplay between a CRC-associated microbe and the extracellular matrix (ECM). Given the importance of the ECM in normal homeostasis and in tumor microenvironment, these findings have important implications in the context of microbial contribution to cancer.

## Introduction

*Streptococcus gallolyticus* subsp. *gallolyticus* (*Sgg*) belongs to the *S. bovis* group of organisms and was previously known as *S. bovis* biotype I [1]. It is an opportunistic pathogen that causes bacteremia and infective endocarditis (IE) [2]. *Sgg* is also known to associate with CRC as documented by numerous case reports and case series over the past several decades [3-7]. A meta-analysis study of case reports and case series published up to 2011 found that among *S. bovis*-infected patients who underwent colonic evaluation, ∼60% had concomitant colon adenomas/carcinomas [8]. Furthermore, patients with *Sgg* bacteremia/IE have a higher risk (∼ 7 fold) for CRC compared to bacteremia/IE caused by other species in the *S. bovis* group [8], suggesting the existence of a *Sgg*-specific mechanism that promotes the strong association between *Sgg* and CRC. The prevalence of *Sgg* in CRC patients has not been investigated as extensively as the risk for CRC among patients with active *Sgg* infections. However, recent data indicate that *Sgg* was enriched in tumor tissues from CRC patients [9-11], suggesting its potential as a biomarker for CRC.

In addition to the strong clinical association between *Sgg* and CRC, studies have shown that *Sgg* stimulates the proliferation of CRC cells and promotes the development of tumors in experimental models of CRC [10, 12-15]. *Sgg* treatment of human CRC cells led to larger tumors compared to untreated cells in a xenograft model. In an azoxymethane (AOM)-induced CRC model, mice orally gavaged with *Sgg* had significantly higher tumor burden and dysplasia grade compared to control mice. In a colitis-associated CRC model, oral gavage of *Sgg* augmented tumorigenesis in the colon. Taken together, long-standing clinical observations and recent functional studies indicate that *Sgg* not only has a strong association with CRC but also actively promotes the development of CRC. The mechanism underlying the tumor-promoting activity of *Sgg*, however, is poorly understood. The ability of *Sgg* to stimulate CRC cell proliferation is an important aspect of the tumor-promoting effect of *Sgg*. The Wnt/β-catenin signaling pathway regulates cell fate and proliferation and is a critical pathway in colon tumorigenesis. Previous results indicated that *Sgg* induced upregulation of β-catenin and increased nuclear translocation of β-catenin, and that β-catenin signaling was required for *Sgg* to stimulate CRC cell proliferation and tumor growth [10]. The signaling events that lead to *Sgg*-induced activation of β-catenin signaling and cell proliferation were unknown.

The extracellular matrix (ECM) regulates fundamental cell behavior such as cell proliferation, adhesion and migration and plays important roles during normal development as well as in pathological conditions such as cancer [16, 17]. The ECM is an important constituent of the tumor microenvironment. Altered ECM composition, structure and mechanical property are common features in tumor tissues and contribute to tumor progression [18-23]. In the case of CRC, multiple studies have found that various types of collagens are upregulated in tumors compared to matched normal tissues [24-31]. Whether gut microbes can provide exogenous signals to modulate ECM expression and dynamics was unknown.

In this study, we found that *Sgg* upregulates the expression of collagen *in vitro* and *in vivo*. We demonstrated that upregulation of collagen by *Sgg* is important for the promotion of CRC cell proliferation, upregulation of β-catenin, and tumor growth by *Sgg*. Moreover, we demonstrated the importance of the ECM in *Sgg*-mediated cancer cell proliferation by using decellularized matrix from CRC cells cultured under various treatment conditions. Altogether, our results suggest a novel mechanism in which *Sgg* actively regulates the expression of ECM molecules which in turn affects the ability of *Sgg* to stimulate CRC cell proliferation in a direct and indirect manner. The results highlight a previously under-studied activity of gut bacteria and have important implications in the context of microbial contribution to CRC.

## Results

### *Sgg* increases collagen expression in human CRC cells and in colonic tissues *in vivo*

*Sgg* was previously shown to stimulate the proliferation of certain human CRC cells including [10, 14]. To investigate the changes in CRC cells induced by *Sgg*, we performed mass spectrometry-based label-free global proteome profiling of whole cell lysates prepared from HT29 cells cultured alone or in the presence of *Sgg* strain TX20005. Strikingly, the level of several types of collagens was increased in cells co-cultured with *Sgg* compared to that in cells cultured alone, with type VI collagen (ColVI) showing the highest relative abundance (supplemental Table S1). The increased expression of ColVI was further confirmed at the transcription and protein level. In RT-qPCR, both ColVI α1 chain (COL6A1) and α3 chain (COL6A3) was significantly increased in the presence of *Sgg* compared to cells cultured in media only (Fig. 1A). In western blot, ColVI level was significantly increased in HT29 and HCT116 cells co-cultured with *Sgg*, compared to cells co-cultured with *L. lactis*, a non-pathogenic negative bacterial control, or in media only (Fig. 1B and 1C). Previous studies showed that *Sgg* stimulates the proliferation of HT29 and HCT116 cells, but had no effect on A549 cells, a human lung cancer cell line [10]. No significant changes in ColVI were observed in A549 cells cultured in the presence of *Sgg* when compared to cells cultured in the presence of *L. lactis* or in media only (Fig. 1B and 1C). Using immunofluorescence (IF) microscopy, we further validated that *Sgg* induced the expression of ColVI (Fig. 1D). Upregulation of type I collagen (ColI) by *Sgg* was also confirmed by using IF (supplemental Fig. S1).

**Fig. 1.**
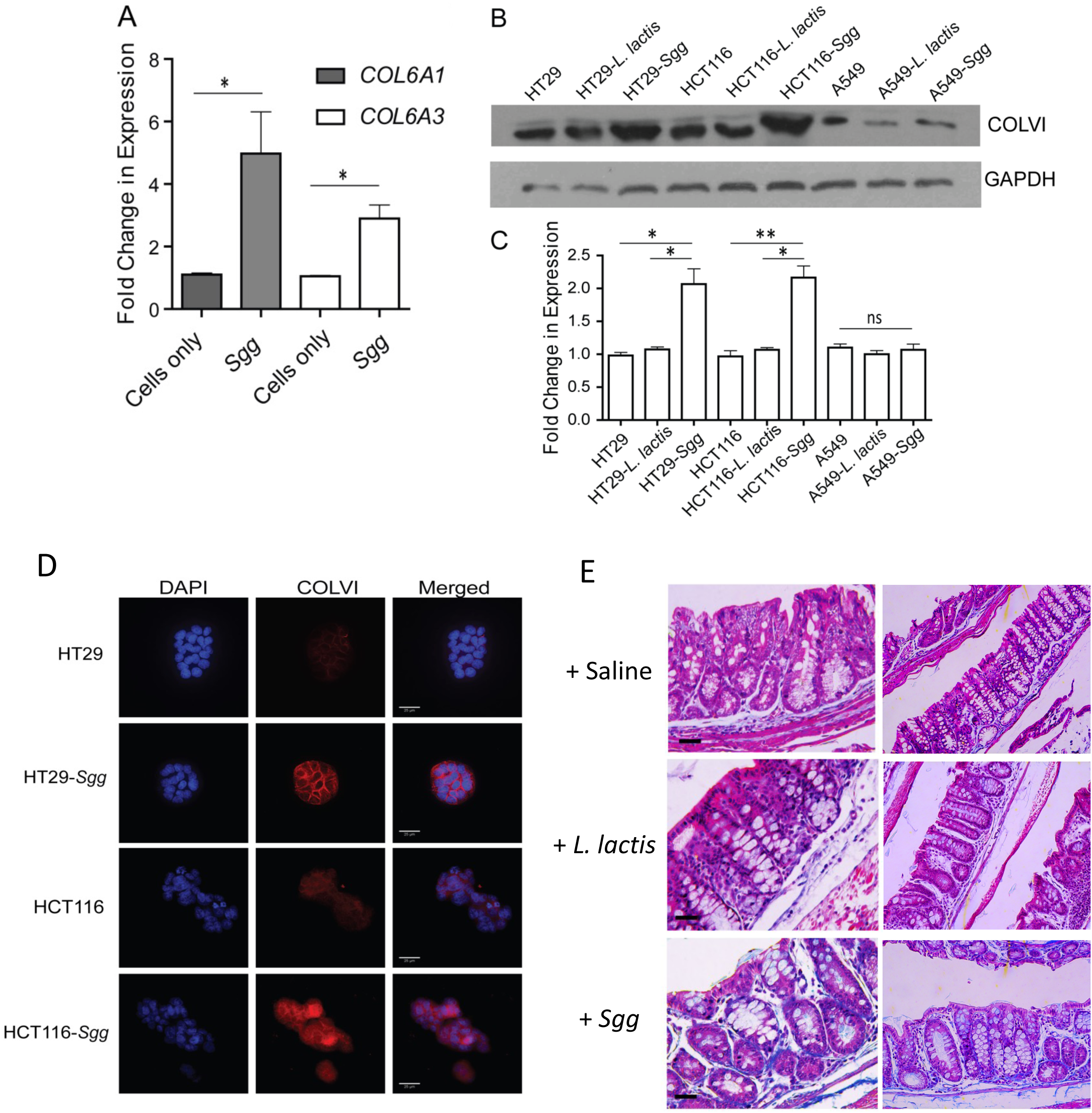
*Sgg* upregulates collagen expression in cultured cells. **A**. HT29 cells were co-cultured with *Sgg* (strain TX20005) or media only for 6 hours. RNA was extracted and analyzed by RT-qPCR. CT values were first normalized to GAPDH then to cells cultured in media only and then converted to fold changes. **B** and **C**. HT29, HCT116 and A549 cells were co-cultured with *Sgg, L. lactis*, or media only for 12 hours. Cell lysates were subject to western blot with anti-ColVI antibody. Band intensity was quantified using Image J and normalized to GAPDH (**C**). **D**. HT29 and HCT116 were co-cultured with *Sgg* or media only for 12 hours. Cells were washed, fixed, incubated with anti-ColVI antibody and counterstained with DAPI. Representative images are shown. Scale bars represent 25µm. **E**. A/J mice were administered with 4 weekly i.p. injections of AOM, followed by treatment with ampicillin for 1 week and then weekly oral gavage of bacteria (*Sgg* and *L. lactis*, respectively) or saline for 12 weeks, as previously described [12, 15]. Colons were harvested one week after the last bacterial gavage, swiss-rolled, fixed with meth-carn, embedded and sectioned. Sections were stained with Trichrome stains. Collagen is stained blue. Statistical analysis in **A** and **C** was done using unpaired, two-tailed *t* test. *, *p* < 0.05; **, *p* < 0.01.

We next examined the effect of *Sgg* on collagen expression *in vivo* using colon sections from mice orally gavaged with *Sgg, L. lactis* or saline. Sections were stained with Masson’s Trichrome stain which stains collagen blue [32]. The results showed that colon sections from mice gavaged with *Sgg* had more intense blue staining compared to sections from mice gavaged with *L. lactis* or saline (Fig. 1E), indicating elevated level of collagen following exposure to *Sgg*. IF staining of the colon sections with an anti-ColVI antibody also showed more intense staining of ColVI in colonic crypts from *Sgg*-gavaged mice compared to control mice (supplemental Fig. S2). Taken together, these results suggest that exposure to *Sgg* results in increased level of collagen in *in vitro* cultured cells and in the intestinal mucosa *in vivo*.

### Collagen is required for *Sgg* to stimulate human CRC cell proliferation

Collagen has been shown to be involved in the proliferation of cancer cells [33-36]. We investigated the role of collagen in *Sgg*’s stimulation of CRC cell proliferation. HT29 COL6A1 and COL6A3 stable knockdown cells were generated. The ability of *Sgg* to stimulate the proliferation of either COL6A1 (Fig. 2A) or COL6A3 (supplemental Fig. S3A) knockdown cells was significantly reduced compared to that in untransfected cells or cells transfected with control shRNA. We confirmed that COL6A1 (Fig. 2B) or COL6A3 (supplemental Fig. S3B) knockdowns reduced the level of ColVI in the cells. *Sgg* was shown to upregulate β-catenin and c-Myc and β-catenin is required for *Sgg* to stimulate cell proliferation [10]. Knockdown of COL6A1 (Fig. 2B-2D) or COL6A3 (supplemental Fig. S3B-S3D) abolished the *Sgg*-induced upregulation of β-catenin or c-Myc, suggesting that ColVI acts upstream of β-catenin in the signaling cascade that leads to *Sgg*-induced cell proliferation. In addition to ColVI, we also carried out knockdown of type I collagen (ColI) using siRNA specific for the α1 chain of ColI (COL1A1). COL1A1 knockdown abolished the ability of *Sgg* to stimulate cell proliferation (supplemental Fig. S4), suggesting that multiple ECM components are involved in *Sgg*-induced cell proliferation.

**Fig. 2.**
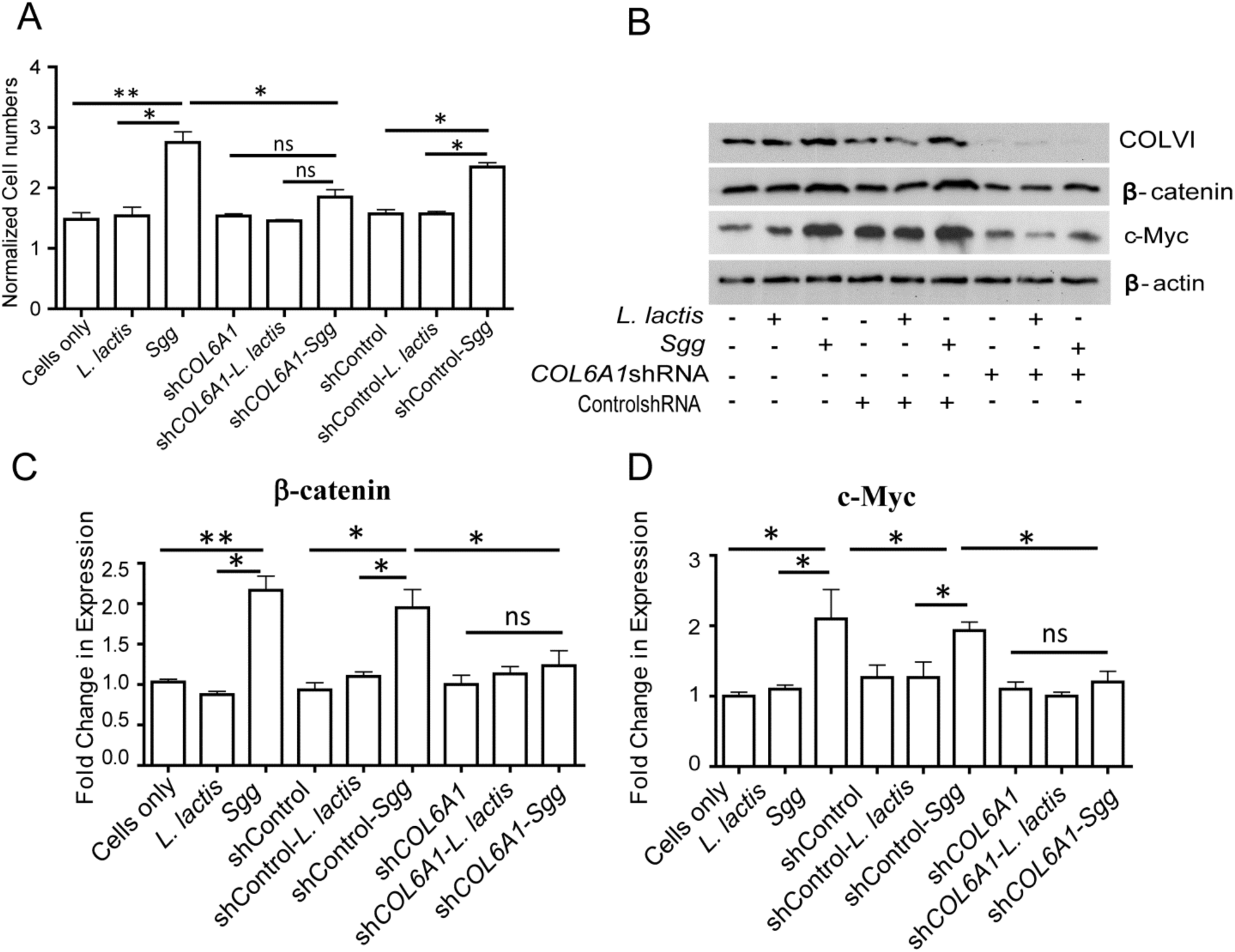
*Sgg* promotes cell proliferation in a ColVI dependent manner. **A**. Knockdown of COL6A1 abolished the effect of *Sgg* on cell proliferation. Untransfected HT29 cells, COL6A1 stable knockdown HT29 cells or HT29 cells transfected with a control shRNA were incubated with media only, *L. lactis* or *Sgg* TX20005 for 24 hours. Cell proliferation assays were performed by counting viable cells as described in the Materials and Methods section. **B-D**. Cells were incubated in media only, *Sgg* or *L. lactis* for 12 hours as described in the Materials and Methods section. Total cell lysates were subject to western blot assays to compare COLVI, β-catenin, and c-Myc protein levels. Representative images are shown (**B**). Band intensity was quantified using Image J, normalized to β-actin first and then to the cells only control (**C-D**). Data are presented as the mean ± SEM. Each experiment was repeated at least three times. Unpaired, two-tailed *t* test was used for statistical analysis. *, *p* < 0.05; **, *p* < 0.01.

### Collagen is involved in *Sgg* adherence to CRC cells

*Sgg* TX20005 was previously shown to bind to several type of collagen including type I, IV and V [37, 38]. We found that TX20005 also attached to immobilized ColVI (supplemental Fig. S5). We investigated if knockdown of ColVI affected the adherence of *Sgg* to CRC cells and found that *Sgg* adherence to COL6A1 knockdown cells was reduced by ∼40% compared to untransfected cells or HT29 cells transfected with the control shRNA (Fig. 3).

**Fig. 3.**
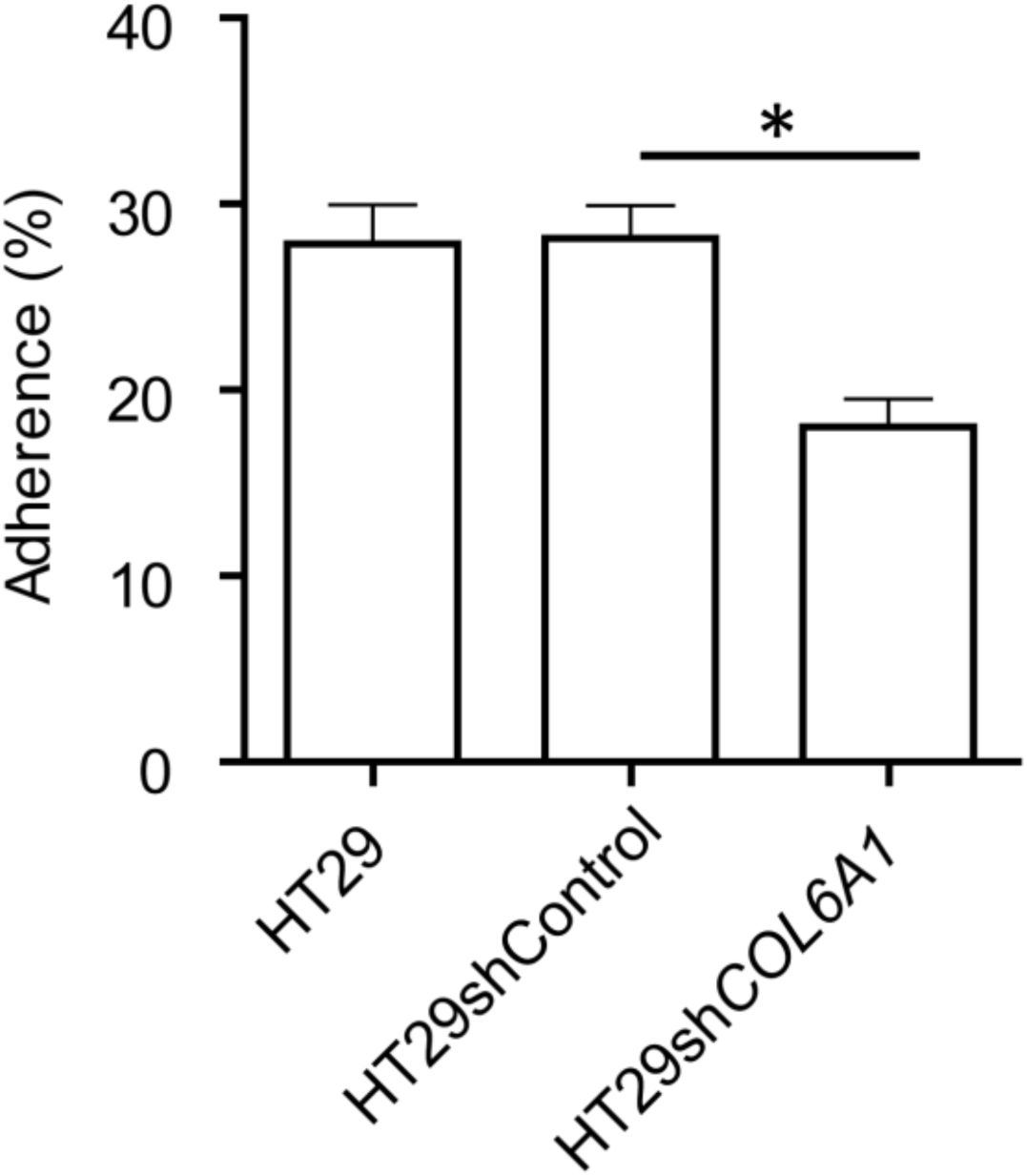
Collagen knockdown impaired the adherence of *Sgg*. HT29 cells, HT29 cells transfected with a control shRNA or COL6A1 stable knockdown HT29 cells were incubated with TX20005 (MOI=10) as described in the Materials and Methods section. Adherence was calculated as the percentage of adhered bacteria vs. total bacterial added and combined from at least three independent experiments. Mean ± SEM is presented. Unpaired, two-tailed *t* test was used for statistical analysis. *, *p* < 0.05.

### Decellularized matrix (dc-matrix) derived from *Sgg*-treated cells alone is sufficient to promote cancer cell proliferation

The importance of collagen in *Sgg-*induced cell proliferation could be a consequence of its effect on *Sgg* adherence to the ECM molecules around CRC cells, leading to a reduced local concentration of *Sgg*. On the other hand, it is known that increased collagen deposition in the matrix leads to matrix-induced cell proliferation [35, 36]. Therefore, it is also possible that collagen contributes to *Sgg*-stimulated cell proliferation in this fashion. To investigate if upregulation of ECM molecules by *Sgg* can directly contribute to cell proliferation, dc-matrix was prepared from HT29 cells cultured in media only, in the presence of *L. lactis* or *Sgg*. We confirmed that no live bacteria were present in the dc-matrix by incubating dc-matrix in antibiotics-free media for 24 hours. No bacterial growth was observed. We also confirmed that no intact cells remained after the decellularization procedure by staining the samples with DAPI (supplemental Fig. S6). HT29 cells were then seeded onto the various dc-matrices and incubated in antibiotics-containing media for 24 hours. The dc-matrix prepared from *Sgg*-treated cells stimulated cell proliferation significantly better than the dc-matrix from HT29 cells alone or from *L. lactis*-treated cells (Fig. 4). Furthermore, dc-matrices were also prepared from COL6A1 knockdown cells that had been incubated with or without *Sgg*. Dc-matrix from *Sgg*-treated COL6A1 cells showed no significant difference in the ability to stimulate cell proliferation compared to dc-matrix prepared from COL6A1 cells alone (Fig. 4). These results indicate that dc-matrix from *Sgg*-treated cells stimulates cell proliferation in a manner that does not require live *Sgg* but depends on collagen. The results also suggest that *Sgg*-induced changes in the ECM play a direct role in stimulating cell proliferation, independent of the effect on *Sgg* adherence.

**Fig. 4.**
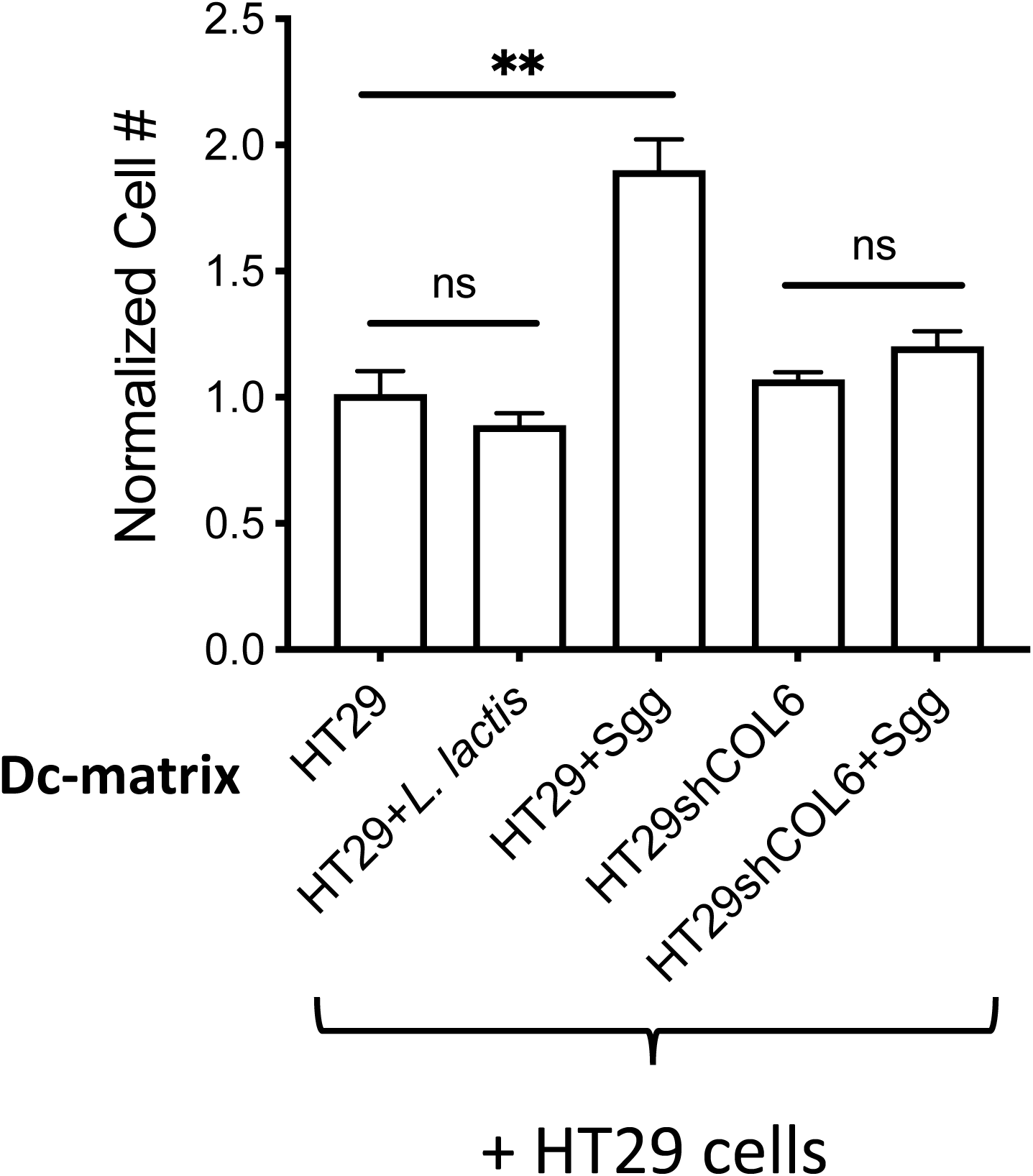
Decellularized matrix (dc-matrix) derived from *Sgg*-treated cells is sufficient to stimulate cell proliferation. HT29 cells were co-cultured with *Sgg* strain TX20005, *L. lactis* or media only for 12 hours. The wells were incubated with antibiotics to eliminate bacteria followed by washing. Cells were then stripped away from the underlying matrix as described in the Materials and Methods section. HT29 cells (∼ 1 × 10^4^) that had not been previously exposed to *Sgg* were seeded on the indicated dc-matrices and incubated for 24 hours. Viable cells were counted. Each experiment was repeated at least three times. Data is presented as the mean ± SEM. Statistical analysis was done using unpaired, two-tailed *t* test. **, *p* < 0.01.

### ColVI is required for *Sgg* to promote tumor growth *in vivo*

To determine the importance of ColVI in *Sgg*-induced tumor growth *in vivo*, shCOL6A1 knockdown cells and cells transfected with control shRNA were cultured in the absence or presence of *Sgg* and then injected into nude mice. For cells transfected with the control shRNA, *Sgg*-treatment resulted in significantly larger tumors at day 7 and 10 post injection compared to untreated cells (Fig. 5). In shCOL6A1 knockdown cells, *Sgg*-treatment did not cause significant increase in tumor size as compared to untreated knockdown cells. We note that in order to prevent infection caused by *Sgg*, mice were administered with antibiotics following injection of cells to eliminate *Sgg*. Therefore, the effect of *Sgg* on tumor growth was more pronounced at the earlier time point than that at the later one. Altogether these results suggest that ColVI is important for *Sgg* to promote tumor growth *in vivo*.

**Fig. 5.**
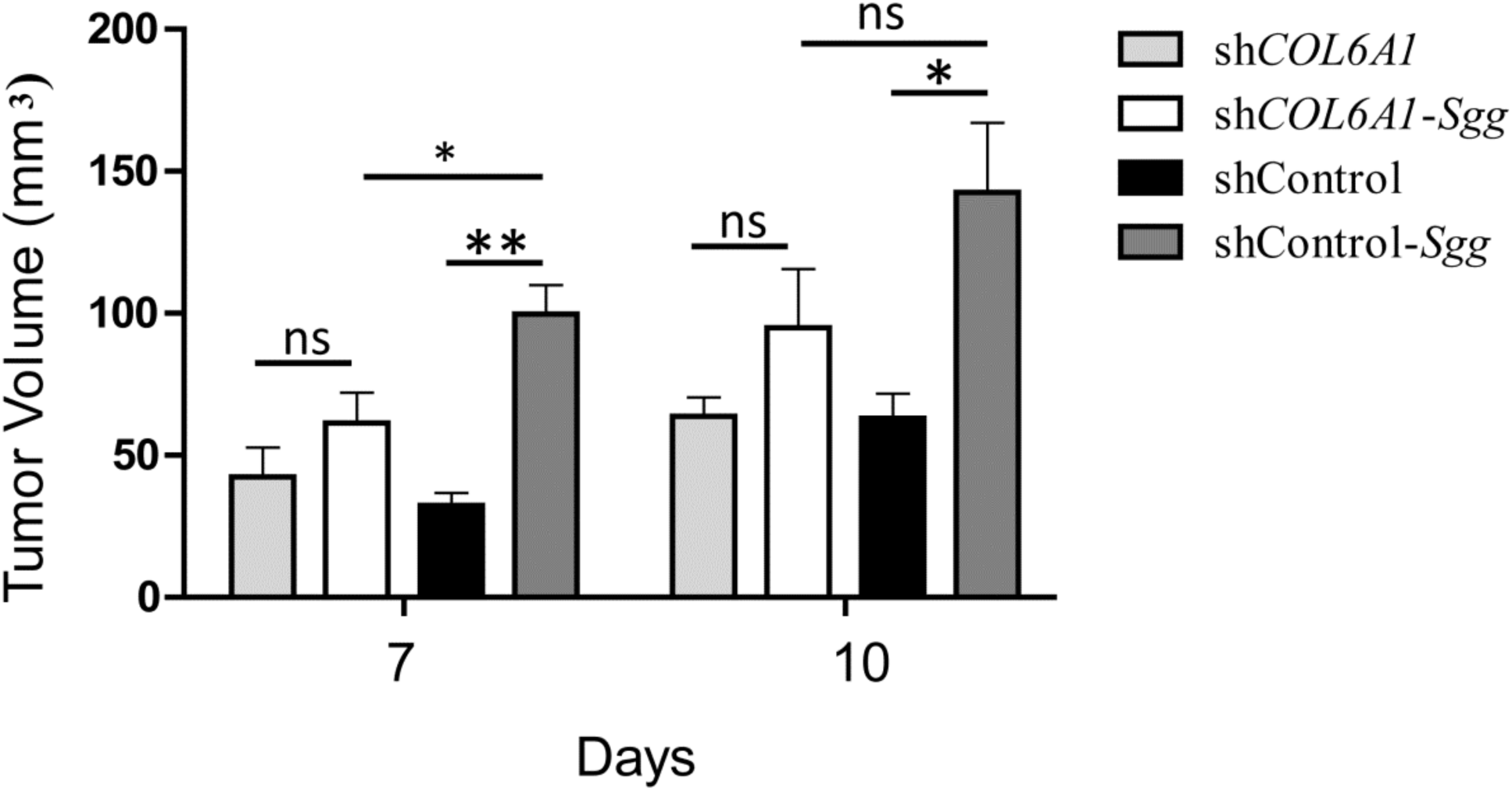
Collagen knockdown impairs the ability of *Sgg* to promote tumor growth *in vivo*. 1 × 10^6^ HT29shCOL6A1 or HT29shcontrol cells were treated with *Sgg* TX20005 or no bacteria, mixed with Matrigel and injected into the dorsal flap of nude mice (n = 5/group). Tumor size was measured at the indicated time point with a digital caliper. Data is presented as the mean ± SEM. Statistical analysis was done using unpaired, two-tailed *t* test. *, *p* < 0.05; **, *p* < 0.01.

## Discussion

It is well appreciated that certain gut microbes or microbial communities play important roles in influencing the development of CRC. Long-standing clinical observations and recent functional studies indicate that *Sgg* is strongly associated with CRC and actively promotes tumor growth, however the molecular mechanisms underlying *Sgg*’s effects are not well understood. The ECM is a component of the tumor microenvironment and an important contributor to tumor progression. However, insights on the impact of gut microbes on the ECM are very limited. Data presented in this study demonstrate that *Sgg* upregulates collagen expression. Furthermore, we provide evidence that *Sgg* upregulation of collagen is important for *Sgg*-induced cell proliferation and tumor growth. To the best of our knowledge, this is the first report that connects a gut microbe with the ECM in the context of microbial contribution to cancer.

Our results showed that *Sgg* upregulates the expression of collagen *in vitro* and *in vivo*. We focused on ColVI since it shows the highest relative abundance revealed by MS analysis. ColVI is also implicated in human CRC in previous studies where its level is higher in tumor tissues compared to matched normal tissues [24-26]. We observed that depletion of COL6A1 or COL6A3 rendered the cells insensitive to the proliferation-promoting effect of *Sgg*. Previous work showed that β-catenin signaling is required for *Sgg*-induced cell proliferation [10]. Results here indicate that upregulation of β-catenin or c-Myc by *Sgg* is abolished in COL6A1 or COL6A3 knockdown cells, suggesting that ColVI is important for *Sgg* to stimulate CRC cell proliferation and acts upstream of β-catenin in the signaling cascade leading to cell proliferation. More importantly, we demonstrated that ColVI is important for *Sgg* to promote tumor growth *in vivo*.

Our results further indicate that collagen contributes to the ability of *Sgg* to stimulate CRC cell proliferation in multiple ways. Previous studies suggested that close proximity of *Sgg* to host cells is important for the pro-proliferative effect of *Sgg* [10, 12]. Knockdown of COL6A1 impaired the adherence of *Sgg*, suggesting that ColVI can influence the ability of *Sgg* to induce cell proliferation by reducing the local concentration of *Sgg*. More importantly, our data also showed that collagen can have a direct effect on *Sgg*-induced cell proliferation. This is supported by results using dc-matrices. Our results showed that dc-matrix prepared from HT29 cells that had been co-cultured with *Sgg* was able to significantly increase cell proliferation compared with dc-matrix from cells cultured alone or with *L. lactis*. On the other hand, dc-matrix from *Sgg* treated COL6A1 knockdown cells showed no difference from that prepared from untreated cells. These results suggest that *Sgg*-modified ECM is directly involved in mediating cell proliferation.

ColVI is special in the sense that a C-terminal cleavage product of COL6A3 chain (endotrophin (ETP)), which is soluble, was found to augment breast tumor growth [39]. Our results using dc-matrix speaks against a role of ETP in increased CRC cell proliferation caused by *Sgg*. ColVI has a unique supramolecular structure among the members of the collagen family. Its beaded microfilament structure enables it to bind to other components of the ECM such as ColI and ColII and acts as a bridging molecule [40-42]. Thus, depletion of ColVI may affect the overall organization and composition of the ECM [17, 43, 44]. In addition, knockdown of ColI also abolished the ability of *Sgg* to stimulate CRC cell proliferation. This, combined with the MS result showing increased expression of several types of collagen in cells co-cultured with *Sgg*, suggests that the effect we observed is not ColVI-specific. It is likely that increased deposition of collagen into the matrix changed the composition of the matrix, resulting in enhanced signaling of collagen receptors. Increased deposition of collagen into the matrix may also affect the mechanical property of the matrix. A stiffer matrix can promote cell proliferation by mechanotransduction [45]. Identifying the signaling pathway(s) downstream of the ECM will shed light on how *Sgg* modification of the ECM contributes to the pro-proliferative effect of *Sgg*.

The molecular mechanisms used by other microbes to drive the development of CRC can be loosely grouped into the following categories: 1) by producing genotoxins that directly induce DNA damage in colonic epithelial cells, 2) by modulating host immune responses to generate a microenvironment favorable for CRC, and 3) by shifting host metabolism to support tumor growth [46-54]. Our results suggest a new way by which microbes influence tumor development - by modulating the ECM. In this study, we focused on the effect of *Sgg* on CRC cells. *In vivo*, fibroblasts are major producers of ECM molecules. Tumor-associated macrophages were also shown to regulate the synthesis and assembly of collagenous matrix [55]. Currently the effect of *Sgg* on fibroblasts and macrophages is unknown. It is possible that *Sgg* may also regulate ECM remodeling by engaging fibroblasts and immune cells *in vivo*.

Components of the ECM such as collagen and fibronectin are commonly targeted by bacterial pathogens to facilitate adherence to and invasion of host cells and colonization of host tissues [56-58]. There is a large body of work investigating the binding interactions between microbial factors and ECM molecules and the biological importance of these interactions. Our results here show that *Sgg* not only targets these ECM molecules for adherence but also actively regulate their expression to provide more attachment sites. More importantly, the *Sgg*-modified ECM is a direct contributor to the pro-proliferative effect of *Sgg*. Thus, our work reveals a novel bidirectional interplay between the pathogen and the ECM. Future work to understand how *Sgg* regulates ECM expression will be important. Given the wide distribution and functional importance of the ECM, modulation of the ECM by *Sgg* is likely to be relevant to *Sgg* pathogenicity in IE. It would also be interesting to see if other bacterial pathogens actively regulate ECM expression to their benefit.

In conclusion, the mechanisms underlying the pro-proliferative and pro-tumor activities of *Sgg* are poorly understood. This study provides the first experimental evidence for *Sgg* modulation of the ECM and a direct role of the ECM in *Sgg*-induced cell proliferation and tumor growth. The results presented here highlight a dynamic two-way interplay between *Sgg* and the ECM and call attention to a novel strategy by which microbes contribute to the development of CRC.

## Materials and Methods

### Bacterial and cell culture conditions

*Sgg* strain TX20005 and *Lactococcus lactis* MG1363 were cultured as described previously [10]. Human colon cancer cell line HT29 and HCT116 were cultured in DMEM/F-12 HEPES (GIBCO, USA) supplemented with 10% fetal bovine serum (FBS) (GIBCO, USA). Human lung carcinoma cell line A549 was maintained in F12-K media supplemented with 10% FBS. All the cells were cultured in a humidified incubation chamber at 37ºC with 5% CO_2_.

### Cell proliferation assays

This was performed as described previously [10]. Briefly, cells (∼1×10^4^ cells/well) were cultured in the presence of stationary phase bacteria (*Sgg* or *L. lactis*) at a multiplicity of infection (MOI) of 1 or media only for 24 hours. Trimethoprim was added at 50 μg/ml final concentration after 6 hours of co-culture to prevent media acidification due to bacterial growth. Cells were detached by trypsin treatment and counted in a Cellometer Mini automated cell counter (Nexcelome Biosciences, Lawrence, MA).

### Collagen knockdown

To generate stable knockdown cells, lentiviral plasmids containing COL6A1 or COL6A3 short hairpin RNA (shRNA) (Sigma-Aldrich, TRCN0000116959 and TRCN0000003622), or MISSION pLKO.1-puro Non-Mammalian shRNA Control (Sigma-Aldrich, SHC002) were first transfected into HEK293T cells to produce lentiviral particles. HT29 cells were then infected with the respective lentiviral particles and selected with puromycin (1μg/ml). Gene knockdown was confirmed by western blot assays. Transient knockdown of COL1A1 was carried out using specific siRNA for COLA1 (ThermoFisher).

### Preparation of decellularized matrix (dc-matrix)

HT29 cells were decellularized following a protocol described previously [59]. Briefly, cells were washed thrice with ice-cold PBS containing a cocktail of protease inhibitors (GenDEPOT). The cells were incubated in a PBS solution containing 0.25% Triton X and 0.25% sodium-deoxycholate in PBS for 5 minutes. The matrix was gently washed in PBS thrice and incubated with 100 mg/mL RNAse A (Roche) and 10 IU/mL DNAse (Sigma) for 30 minutes followed by three washes in PBS.

### Quantitative reverse transcription PCR (RT-qPCR)

HT29 cells were co-cultured with *Sgg* for 6 hrs. Total RNA was extracted from co-cultured cells using the RNeasy Kit (QIAGEN). cDNA was generated by using the Transcriptor First Strand cDNA Synthesis Kit (Roche). qPCR was performed using FastStart SYBR green master mix (Roche) in a Viia 7 Real Time PCR System (Applied Biosystems). The following cycle conditions were used: 95°C for 10 minutes followed by 40 cycles at 95°C for 30 seconds and 60°C for 1 minute. CT values were first normalized to GAPDH then to cells cultured in media only.

### Western blot assays

This was performed as described previously [10]. Briefly, cells were cultured in the appropriate medium in the presence or absence of bacteria for 12 hours. Cells were washed with sterile PBS three times and lysed. Total cell lysates were subjected to SDS-gel electrophoresis and western blot. Rabbit polyclonal antibodies against ColVI (1:500, Abcam), β-catenin (1:4000, Cell Signaling Technology (CST)), c-Myc (1:3000, Abcam), and β-actin (1:5000, CST) were used. Horse radish peroxidase (HRP)-conjugated anti-rabbit IgG (CST) was used as the secondary antibody. Signals were detected using HyGLO, chemiluminescent HRP (Denville, Mteuchen, NJ). Band intensity was quantified using Image J.

### Adherence Assay

This was performed as described previously [10]. Briefly, HT29 cells were incubated with or without bacteria at an MOI of 10 for 1 hour. The wells were washed three times with sterile PBS to remove unbound bacteria. To determine the number of bound bacteria, cells were lysed with sterile PBS containing 0.025% Triton X-100 and dilution plated. Adherence was expressed as a percentage of total bacteria added.

### Animal experiment

Animal studies were performed in accordance with protocols approved by the Institutional Animal Care and Use Committee at the Texas A&M Health Science Center, Institute of Biosciences and Technology. The xenograft model experiment was performed as described previously [10]. Tumor diameters were measured with a digital caliper, and tumor volume calculated using the formula: Volume = (d1xd1xd2)/ 2, with d1 being the larger dimension.

### Immunofluorescence

**1) Colon sections.** Methcarn-fixed paraffin embedded colon sections were deparaffinized with xylene and rehydrated in an ethanol gradient. The slides were incubated in a citrate buffer at 95ºC for 15 minutes, cooled to room temperature (RT), rinsed with PBS and incubated in blocking buffer (PBS containing 1% Saponin and 20% BSA) for 30 minutes. The slides were then incubated with rabbit anti-ColVI (1:200, Abcam) at 4ºC overnight, washed with PBS, and incubated with donkey-anti-rabbit Alexa 594 for 1 hour at RT. The slides were washed again, stained with DAPI, mounted and examined in a DeltaVision Elite microscope (GE Healthcare). **2) Cultured cells**. Cells were seeded onto an 8-chambered slide and cultured under various conditions. Cells were washed 3 times in PBS to remove unbound bacteria, fixed with 4% formaldehyde, and permeabilized with 0.1% Triton-X-100 for 30 minutes. Cells were then incubated in a blocking solution (PBS containing 5% donkey serum and 0.3% Triton X-100) for 1 hour. The slides were then incubated with anti-Collagen VI or anti-Collagen I antibody (1:100) at 4ºC overnight, washed with PBS, and incubated with donkey-anti-rabbit Alexa 594 (1:500 dilution in PBS) for 1 hour at RT. The slides were washed again, stained with DAPI, mounted and examined in a DeltaVision Elite microscope (GE Healthcare).

### Trichrome staining of colon sections

Colon sections were deparaffinized, rehydrated and stained using a Trichrome Stain Kit (Abcam, ab150686) following the protocol provided by the manufacturer.

### Ethics statement

Animal studies were performed in accordance with protocols (IACUC#2017-0420-IBT) approved by the Institutional Animal Care and Use Committee at the Texas A&M Health Science Center, Institute of Biosciences and Technology. The Texas A&M University Health Science Center—Institute of Biosciences and Technology is registered with the Office of Laboratory Animal Welfare per Assurance A4012-01. It is guided by the PHS Policy on Human Care and Use of Laboratory Animals (Policy), as well as all applicable provisions of the Animal Welfare Act. Mice were euthanized by CO_2_ inhalation followed by cervical dislocation.

## Acknowledgements

This work was supported by funds from the Hamill Foundation and the Cancer Prevention and Research Institute of Texas (CPRIT) (RP170653). BCM Mass Spectrometry Proteomics Core is supported by the Dan L. Duncan Comprehensive Cancer Center NIH award (P30 CA125123), CPRIT Core Facility Award (RP210227)

## Figure Legends

**Supplemental Table S1.**
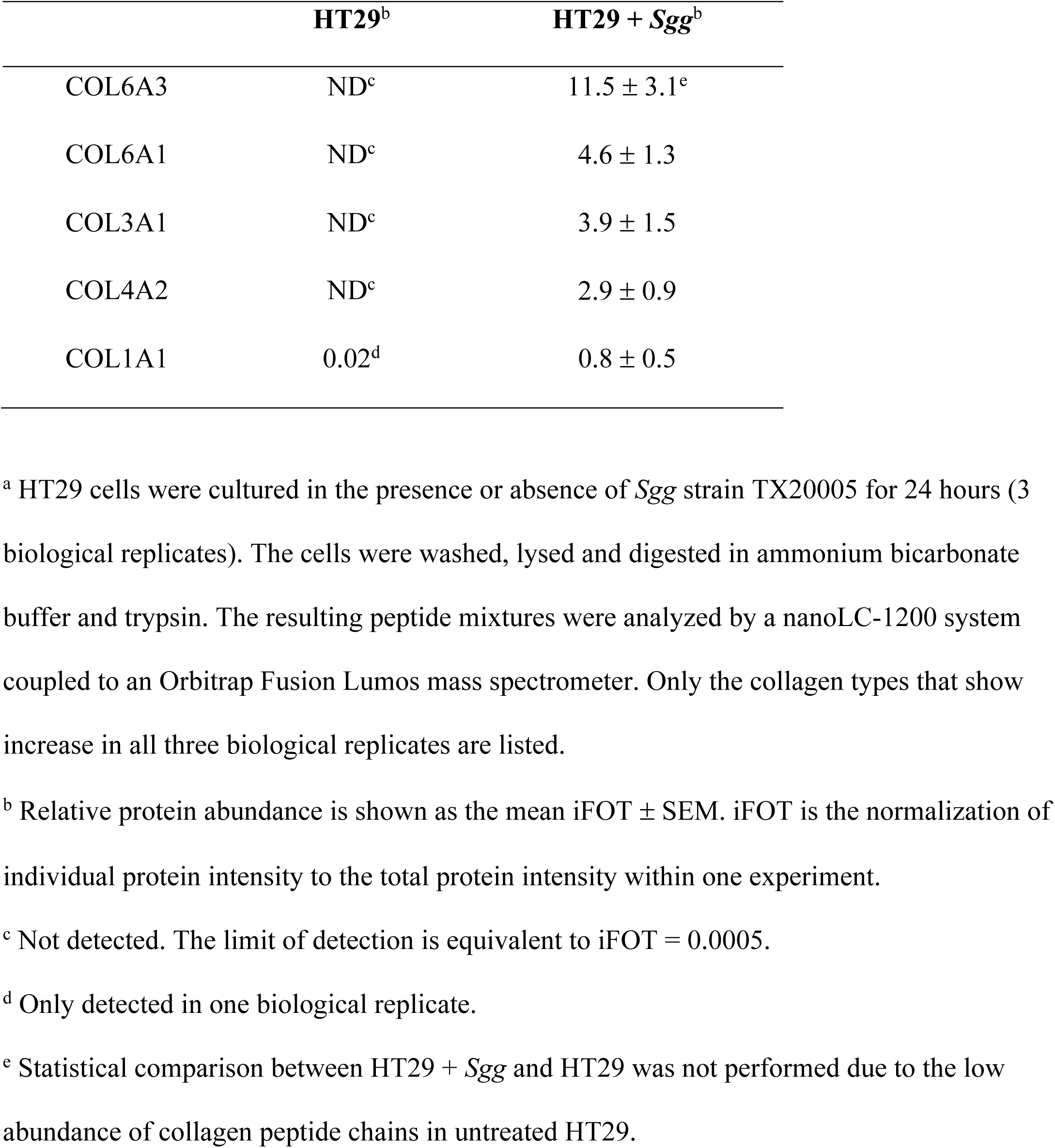
Relative abundance of several types of collagens in whole cell lysates analyzed by Mass Spectrometry ^a^.

---

**Supplemental Fig. S1.**
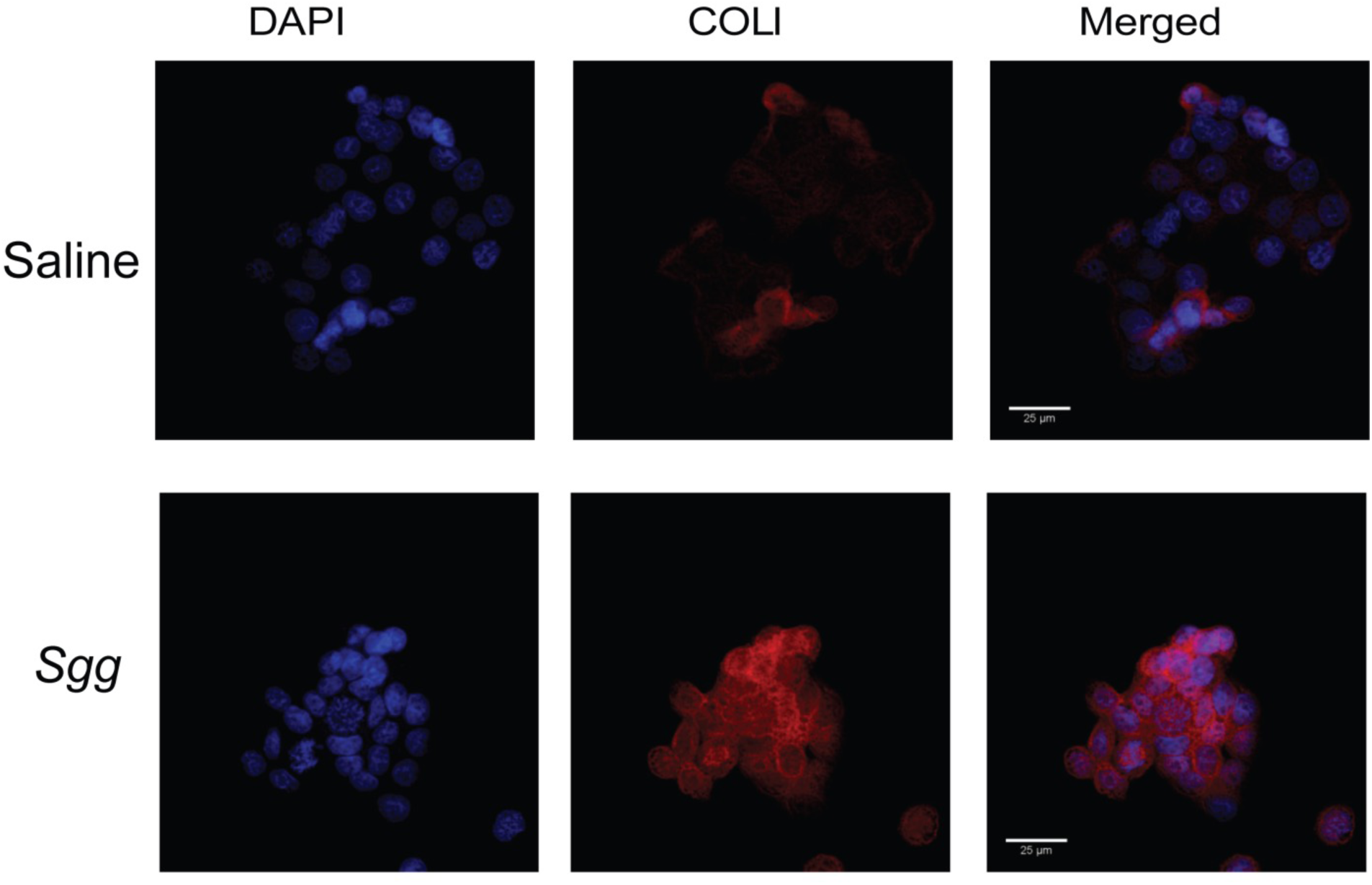
*Sgg* upregulates type I collagen. HT29 cells were co-cultured with *Sgg* TX20005 or media only for 12 hours. Cells were washed, fixed, incubated with anti-ColI antibody and counterstained with DAPI.

**Supplemental Fig. S2.**
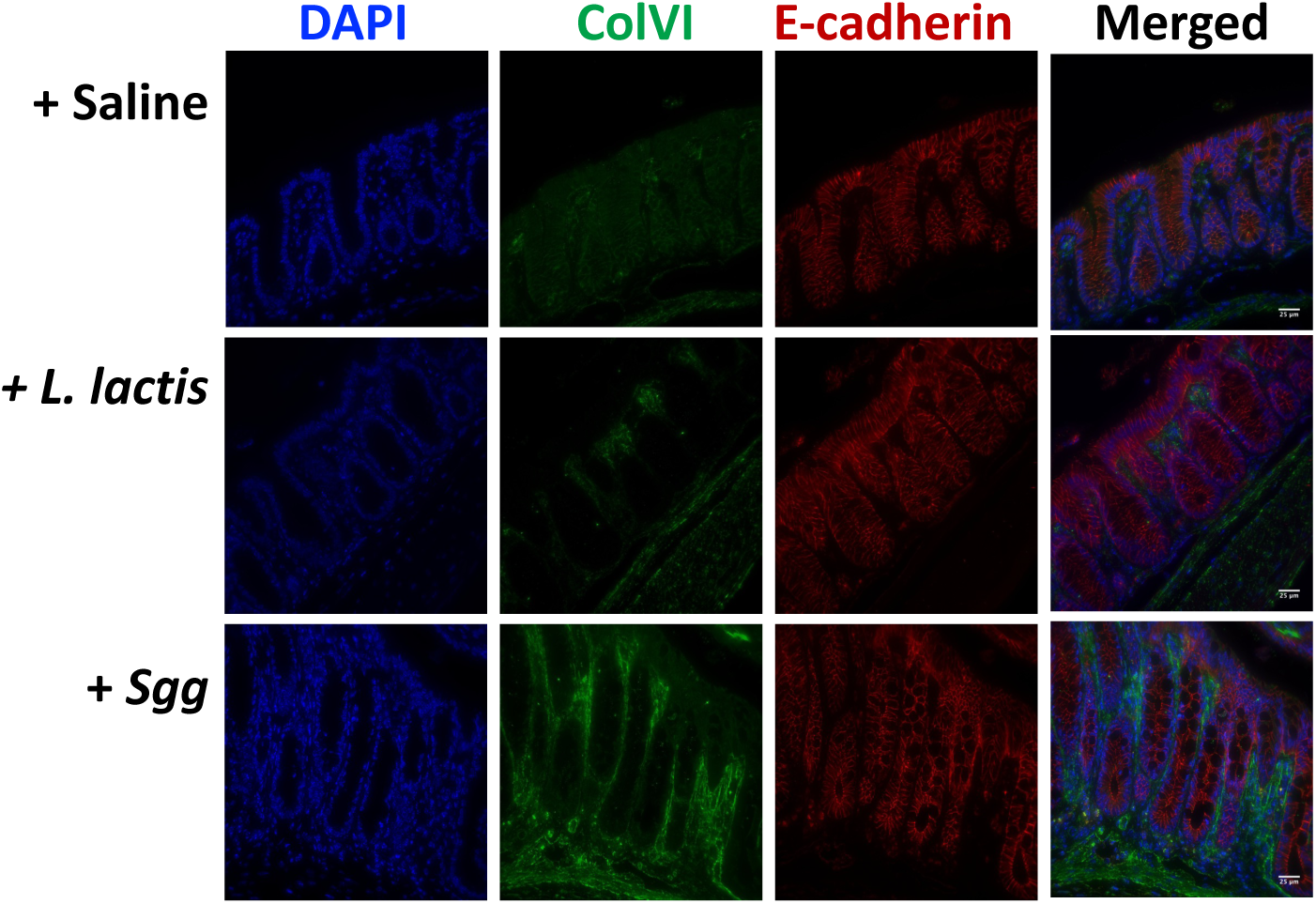
*Sgg* upregulates type VI collagen *in vivo*. A/J mice were administered with 4 weekly i.p. injections of AOM, followed by treatment with ampicillin for 1 week and then weekly oral gavage of bacteria (*Sgg* and *L. lactis*, respectively) or saline for 12 weeks. Colons were harvested one week after the last bacterial gavage, swiss-rolled, fixed with meth-carn, embedded and sectioned. Colon sections were incubated with antibodies against COLVI and E-cadherin (to indicate colonic epithelial cells), followed by appropriate secondary antibodies and counterstained with DAPI.

**Supplemental Fig. S3.**
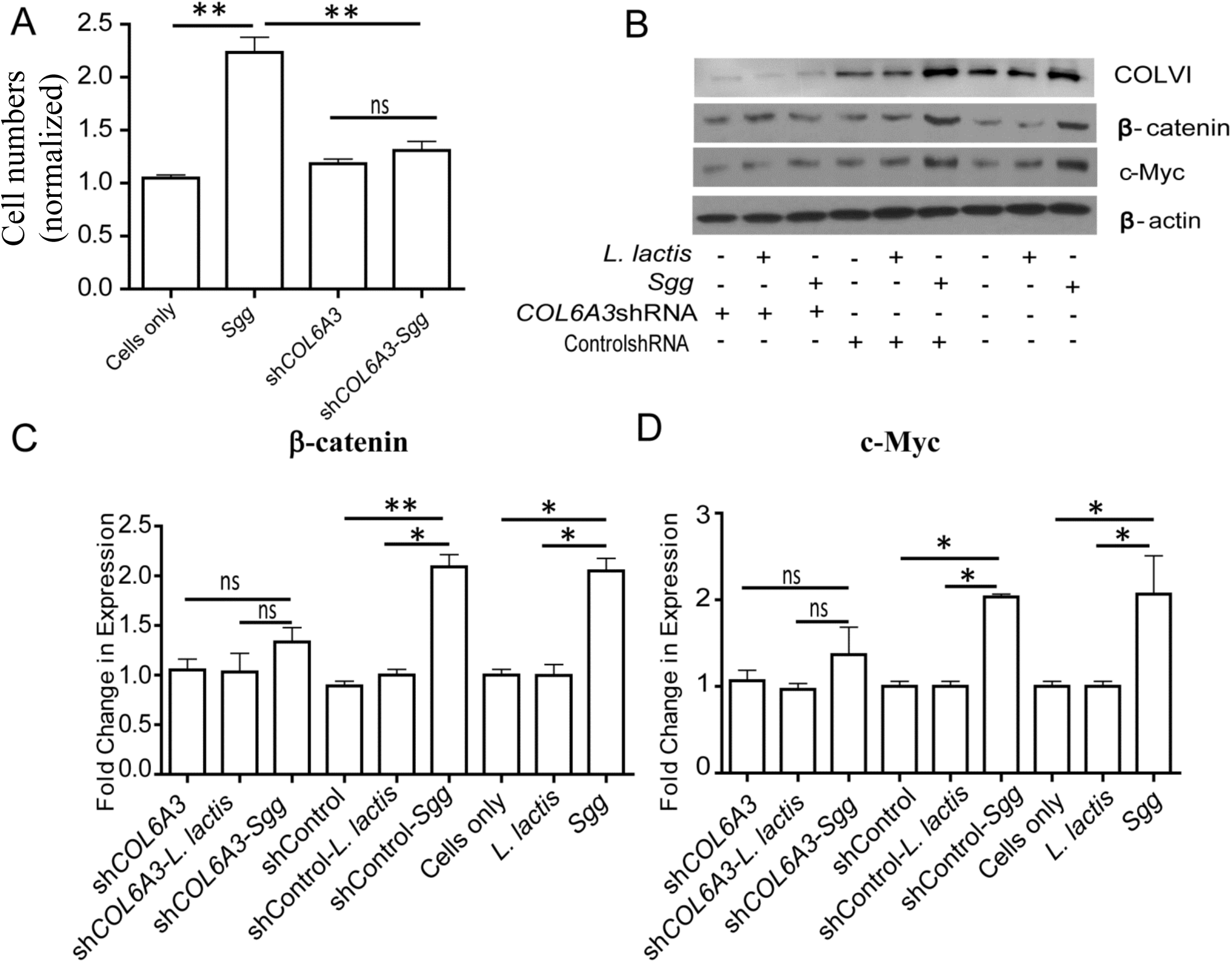
Knockdown of Col6A3 rendered *Sgg* unable to stimulate cell proliferation. The experiments were carried out as described in Materials and Methods section. **A. Cell proliferation assay**. HT29 cells and COL6A3 stable knockdown HT29 cells were incubated with *Sgg* TX20005 or media only for 24 hours. Viable cells were counted using an automated cell counter. **B-D. Western blot** of whole cell lysates from HT29 cells, HT29 cells with control shRNA, or COL6A3 stable knockdown HT29 cells incubated with *Sgg* TX20005 or *L. lactis* for 12 hours. Band intensity from three independent experiments were quantified and normalized to β–actin. Fold change was against shCOL6A3 cells incubated in media only. Unpaired two tailed t test was used for pairwise comparison. *, *p* < 0.05; **, *p* < 0.01; ***; ns, not significant.

**Supplemental Fig. S4.**
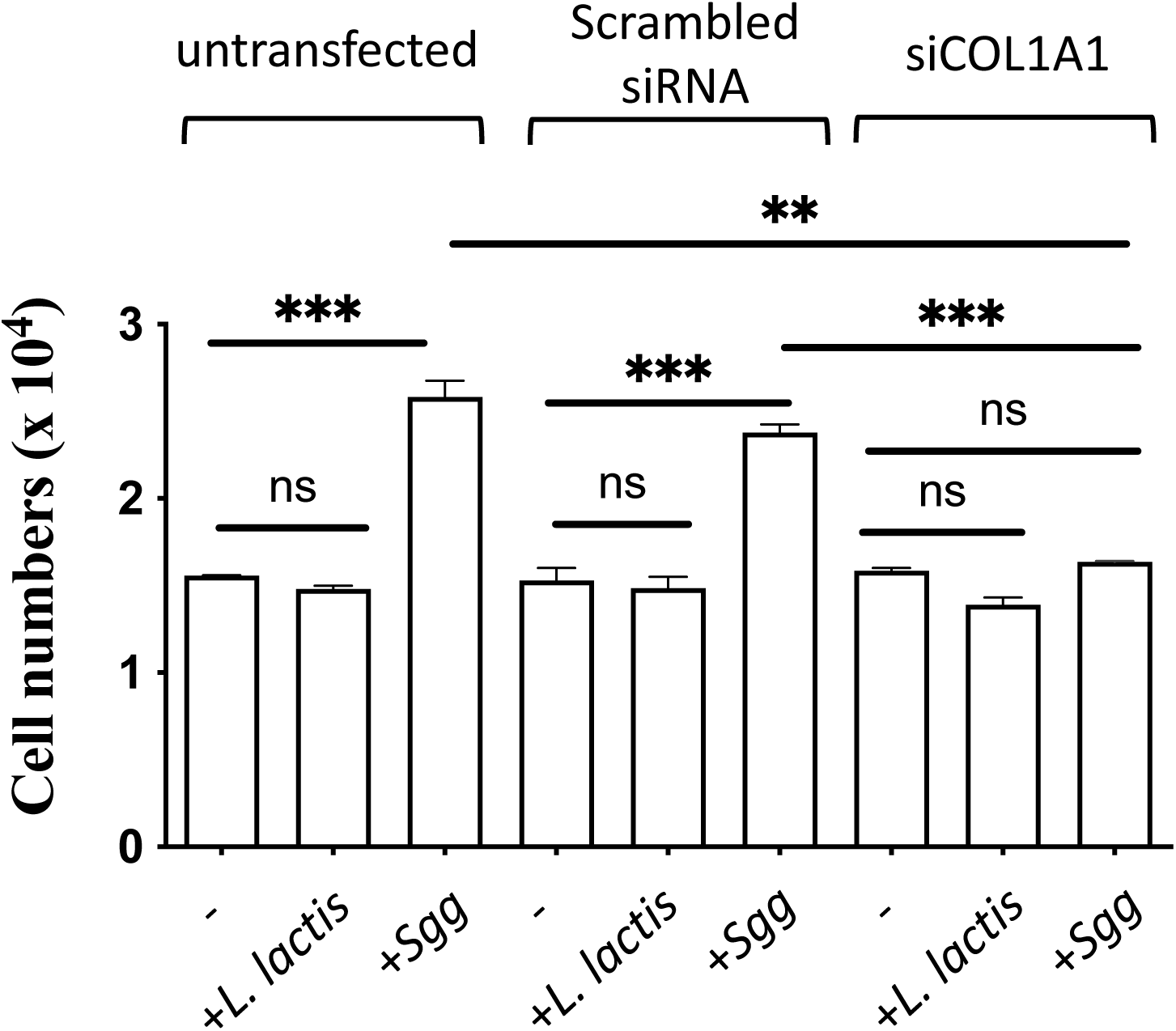
Knockdown of Col1A1 rendered *Sgg* unable to stimulate HT29 cell proliferation. Cell proliferation assay was carried out as described in the Materials and Methods section. HT29 cells were transfected with siRNA for COL1A1 or scrambled control siRNA and incubated for 24 hours. The cells were then incubated with *L. lactis* or *Sgg* TX20005 for 24 hours. Viable cells were enumerated using an automated cell counter.

**Supplemental Fig. S5.**
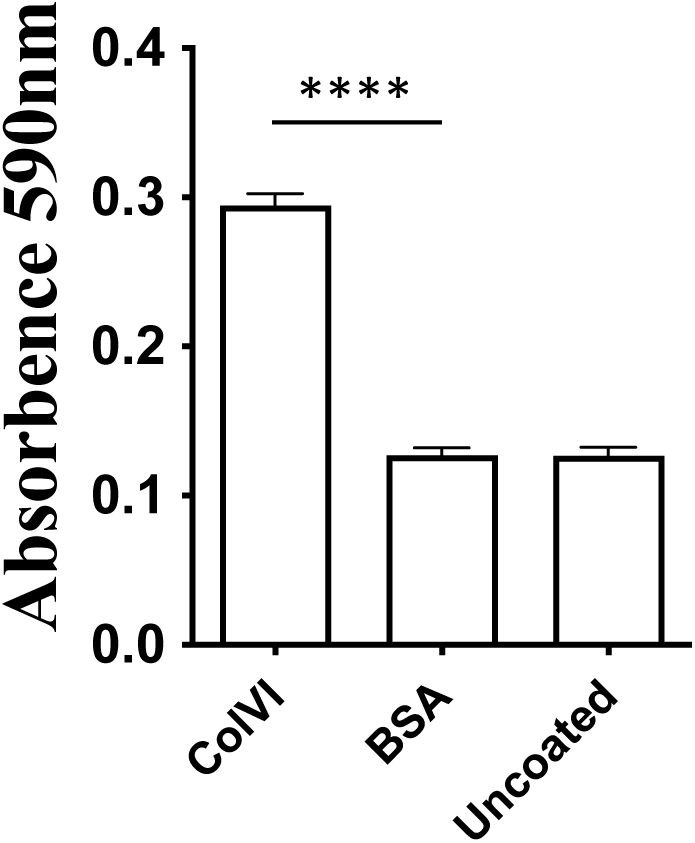
*Sgg* binds to immobilized ColVI in a dose-dependent and saturable manner. ColVI or BSA were coated into 96-well plate at 1µg/well and blocked. Uncoated wells were incubated with PBS. Overnight cultures resuspended in PBS were then added to the immobilized proteins and incubated for 1 hour at room temperature. The wells were washed three times with PBS. Bound bacteria were detected by using the crystal violet staining method.

**Supplemental Fig. S6.**
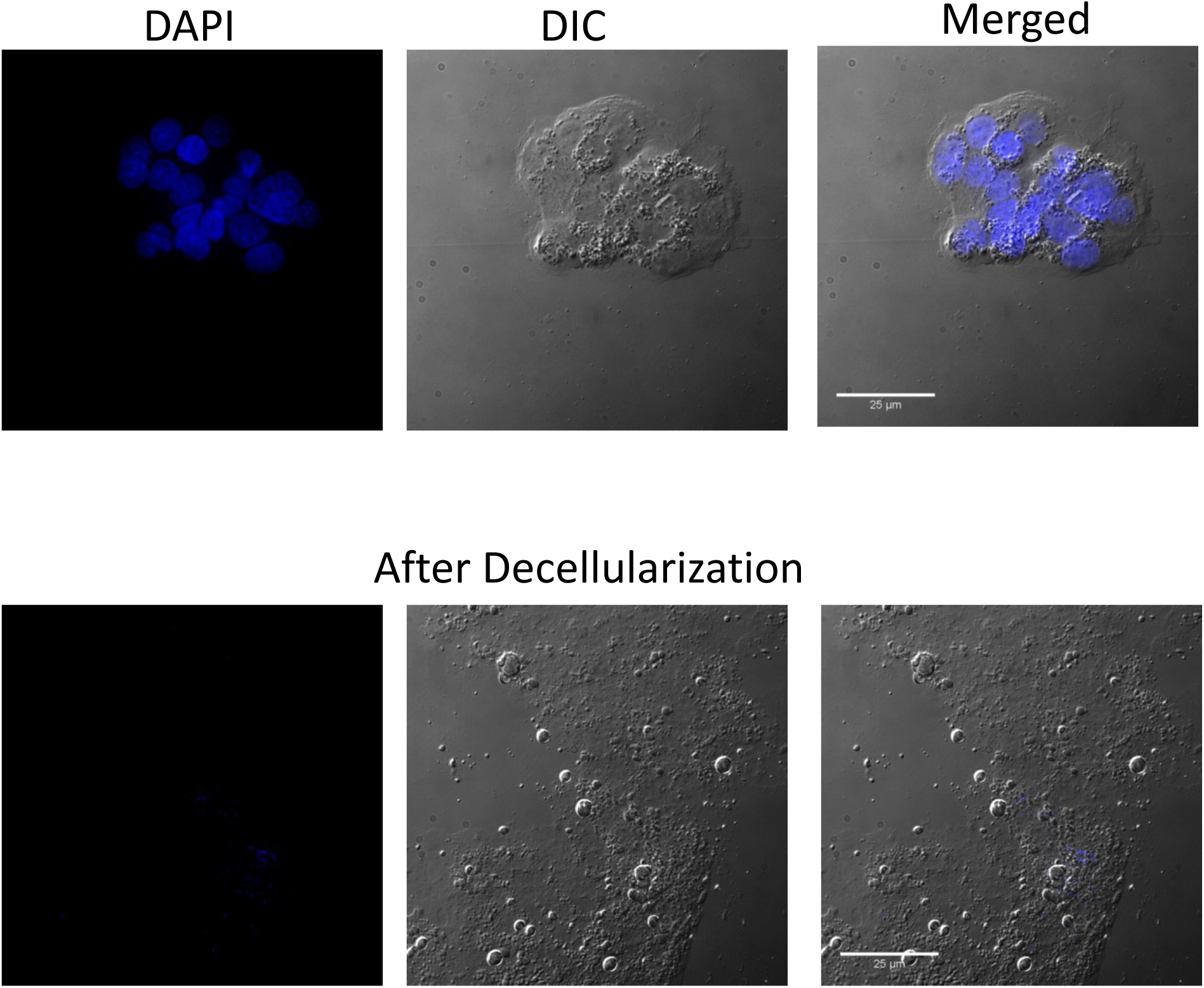
Decellularization of HT29 cells. HT29 cells were treated with 0.25% Triton-X and 0.25% Sodium-Deoxycholate for 5 minutes and then with DNAse and RNAse for 30 minutes, as described in Materials and Methods. The samples were washed thrice with PBS and then fixed and stained with DAPI.

